# Phylodynamics and genome-wide association studies reveal the expansion of modern *Streptococcus canis* in Germany

**DOI:** 10.1101/2025.09.29.679251

**Authors:** Etienne Aubry, Andries J. van Tonder, Miriam M.D. Katsburg, Hayley J. Wilson, Antje-Maria Lapschies, Geoffrey Foster, Stefan Schwarz, Torsten Semmler, Antina Lübke-Becker, Andrew Waller, Julian Parkhill, Marcus Fulde

## Abstract

*Streptoccocus canis* is a leading canine pathogen causing 22.4% of canine streptococcal infections. However, knowledge of factors contributing to *S. canis* expansion is limited. This study uses population genetics to structure a *S. canis* dataset of 585 isolates and identify genetic markers contributing to the success of each dominant lineage. The dataset is composed of canine, feline and bovine isolates primarily from Germany collected between the years 1996 and 2021. We performed Multi-Locus Sequence Typing (MLST) for an initial analysis and clustered the population with *fastBAPS* based on the whole genome. Dated phylogenies were inferred with *BEAST* and Genome-Wide Association Studies (GWAS) were conducted with *Scoary*. MLST showed that in Germany there are two dominant groups of canine *S. canis*, ST43 (n= 75) and ST9 (n= 51), which were grouped into BAPS-6 and BAPS-5, respectively. The BAPS-6 cluster emerged as early as 1988 with major expansion starting around 2010. We showed via GWAS that this cluster is associated with a putative *Streptococcus anginosus* derived integrative and conjugative element carrying a putative cysteine protease. Furthermore, the BAPS-5 cluster emerged around 1786 with a bovine/feline subcluster appearing around 1879. This subcluster is associated with a variant of the streptococcal *lac* operon, which appears to have resulted from an exchange with *Streptococcus dysgalactiae.* We present the largest population genetics study to date for *S. canis* where we show that its expansion is associated with genetic exchange with other streptococcal species leading to increased pathogenic potential (BAPS-6) or host adaptation (BAPS-5).

**Importance:** *Streptococcus canis* is a bacterium which poses considerable threat to the health of dogs through superficial infections. Current research could benefit from understanding the reasons for its expansion and spread into other hosts. We looked at the genomes of 585 *S. canis* isolates collected during a 25-year period and identified which parts of the population were the most successful. With powerful statistical methods, we found that two groups of *S. canis* were the most effective: a canine group which had become more pathogenic by acquiring DNA from *Streptococcus anginosus*, and another group which spread from dogs into cats and cows by acquiring genetic material for lactose consumption from *Streptococcus dysgalactiae*. For the first time, we link the expansion of major *S. canis* groups to genetic exchange events with closely related streptococcal species to understand *S. canis* evolution and spread.

## Introduction

*Streptococcus canis* is a major Lancefield group G β-haemolytic opportunistic pathogen which has recently been discovered to also express Lancefield group C antigens (1). It is responsible for 22.4% of streptococcal canine infections and is typically identified as a commensal bacterium in dogs and cats, but in cases of infection it is known for its ability to cause keratitis and otitis externa (1–4). It has also been reported to cause more severe systemic infections, such as streptococcal toxic shock-like syndrome, necrotising fasciitis, and infective endocarditis (5–7). Recent research has identified *S. canis* as a multi-host and zoonotic pathogen capable of infecting cattle, seals, rats, and mink (2, 8–12). The multi-host nature of *S*. *canis* has raised questions about its transmission routes which could lead to further infections (13). Of note is the cross-infection between feline and bovine hosts where feline carriers of *S. canis* are suggested to be the most likely source of certain bovine mastitis outbreaks (14). This was previously shown during an outbreak that was caused by *S. canis* ST55, which was recovered from both cattle and a cat in the same farm (15).

Despite the veterinary relevance of *S. canis,* knowledge of its pathogenesis is still lacking with the best characterised virulence factor being the *S. canis* M protein (SCM). The M protein of *S. pyogenes* is a critical and highly versatile virulence factor and is one of the leading candidates for a vaccine against this major human pathogen (16, 17). It has been shown that SCM is equally as versatile, with previous studies observing its capacity to bind (mini)-plasminogen, IgG, and fibrinogen (18–20). The arginine deiminase system is the only other characterised genetic maker which is considered to be implicated in *S. canis* pathogenesis (21). Many other virulence factors have been identified in notably *S. pyogenes* which could be interesting targets for *S. canis* pathogenesis, as well. Cysteine proteases, such as SpeB, have gained great attention in research due to their broad spctrum of functions (22). Indeed, SpeB has been shown to degrade over 200 host proteins, including immunoglobulin, fibrinogen and plasminogen, thereby aiding in host immune evasion. *SpeB* was even proven to be regulated by the LacD component of the streptococcal *lac* operon where it was then postulated to be a regulator of *S. pyogenes* virulence in the presence of appropriate nutritional conditions (23).

The rising accessibility of whole genome sequencing (WGS) has allowed epidemiological studies to provide new insights into pathogens on a population level. Population genetics studies have become essential to help understanding the expansion and evolution of many pathogens (24). Notably, one major study on the zoonotic pathogen, *Streptococcus suis,* has linked the expansion of pathogenic lineages to the acquisition of three pathogenicity islands (25). Host adaptation signals have been identified in *Streptococcus agalactiae* where human isolates were linked to the presence of *scpB,* bovine isolates to the *lac* operon and piscine isolates to the novel Locus 3 (26). Similar studies have been conducted on *S. canis* where it was shown through Multi-Locus Sequence Typing (MLST) and core genome analysis that *S. canis* is dominated by Sequence Type 9 (ST9) and has the potential for zoonotic transfer (2, 11, 27). Complementing these studies with methods that provide higher levels of resolution and larger sampling sizes would further clarify the evolutionary pathway of *S. canis*, which could identify and explain the expansion of pathogenic lineages.

This study, therefore, aims at characterising the population structure of the largest *S. canis* dataset collected to date to identify its major lineages and to understand the possible mechanisms driving their expansion. Earlier research has shown that *S. pyogenes, S. agalactiae* and *S. dysgalactiae* subsp. *equisimilis* have expanded due to the acquisition of novel virulence factors or antimicrobial resistance genes through horizontal gene transfer (28–30). *S. canis* has been shown to be involved in genetic exchange with its close relative *S. dysgalactiae* subsp*. equisimilis* with the newly identified Lancefield C *S. canis* possibly arising from one of these exchanges (1, 27). The initial characterisation of the genome of *S. canis* showed that much of its recent evolution could be attributed to the acquisition of mobile genetic elements carrying virulence factors (31). Of note, genetic exchanges with the bovine pathogens, *S. agalactiae* and *S. dysgalactiae* subsp*. dysgalactiae* were detected providing possible opportunities for host adaptation (31). We hypothesize that the major *S. canis* lineages could be expanding due to the acquisition of virulence factors or antimicrobial resistance genes via horizontal gene transfer from other streptococcal species.

Here we use Bayesian Analysis of the Population Structure (BAPS) to show that a 585-sample population of *S. canis* can be structured into 12 distinct BAPS clusters and that in Germany between the years 1996 and 2020, the main dominant lineage of *S. canis* has shifted from BAPS-5 to BAPS-6. Through ancestral state reconstruction and Genome-Wide Association Studies (GWAS), we show that the emergence of the new BAPS-6 cluster is associated with the acquisition of a putative Integrative and Conjugative Element (ICE) from *S. anginosus.* We also show that BAPS-5, the largest cluster in the dataset, presents bovine and feline host adaptation signals associated with the acquisition of a *lac* operon variant obtained from *S. dysgalactiae*.

## Results

### General population structure of *S. canis*

We sequenced a collection of 468 isolates (36 feline isolates, 68 bovine, 360 canine, and four unknown) and combined them with 117 publicly available genomes (16 feline isolates, 48 canine, 25 human, two pinniped isolates, and 26 unknown) from the European Nucleotide Archive (ENA) to produce a total dataset of 585 whole genome sequences. Isolates were recovered from almost every region in Germany with the highest sampling densities corresponding to the Berlin region (n= 126), followed by Bavaria (n= 114) and Lower Saxony (n= 48) (**Fig 1a**). We conducted an initial analysis of this population by evaluating the isolates based on Multi-Locus Sequence Typing (MLST) to compare them to the existing Pinho dataset which showed that *S. canis* was dominated by ST9 (11, 27). We found that our collection was also dominated by ST9 (accounting for 21.7% of the entire test population) but this was closely followed by the so far underreported ST43 population (15.4%) with the next most frequently represented ST being ST13 (4.5%). When plotting the proportions of the German canine *S. canis* population between the years 1996 to 2020, we found that, between 1996 and 2015, ST9 was the main representative of the population accounting for more than 25% of the dataset at the time (n= 21). However, from 2016 on, we detected ST43 in the dataset which then dominated the population representing a more than 25% prevalence (n= 76). In contrast, ST9 decreased to less than 10% prevalence (n= 33) during this time (**Fig 1b**). The canine isolates which were present in the rest of the world showed that from 2002 to 2015, ST9 accounted for more than 25% of the isolates (n= 4) (**Fig 1c**). The prevalence of ST9 decreased to 15% of the isolates during the 2016-2021 period (n= 12) and ST43 was detected to account for 7.5% (n= 6).

**Fig 1:**
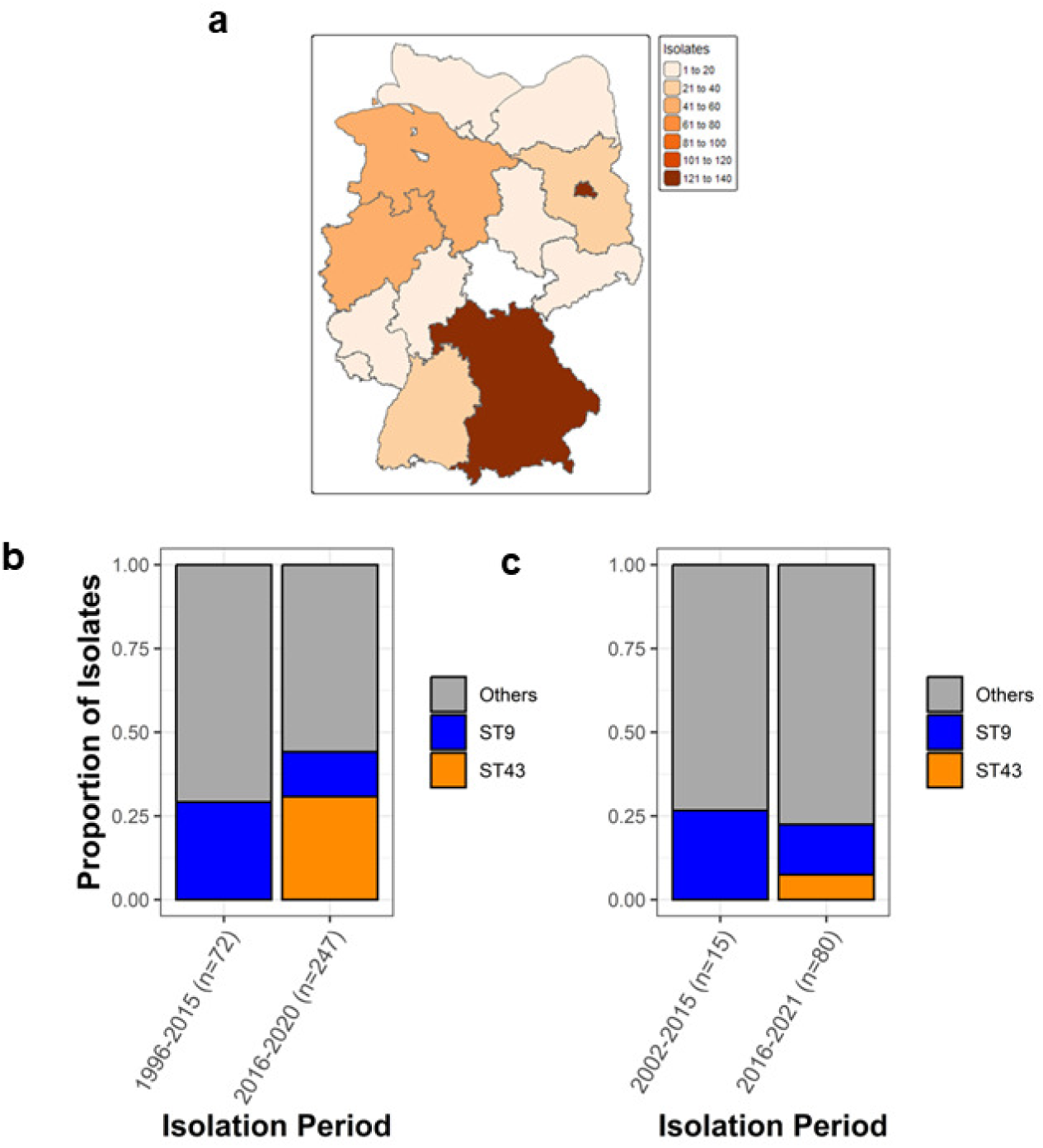
Epidemiological data of *S. canis* worldwide and in Germany. Epidemiological map of the 458 German *S. canis* isolates. A darker colour denotes higher sampling density (a). Stacked barplot representing the proportions of dominant *S. canis* Sequence Types in German canine hosts between the years 1996 and 2020 (b). Stacked barplot representing the proportions of Sequence Types found in canine hosts outside of Germany between the years 2002 and 2021 (c). ST9 and ST43 are represented in blue and orange. Grey denotes all other sequence type population.

To examine the population change with a higher resolution, we analysed the population structure with *fastBAPS* based on whole genome alignments mapped to the reference sequence HL_77. We found that the population was partitioned into 12 distinct BAPS clusters. We detected BAPS-5 (n= 183, 31.3% of dataset), BAPS-6 (n= 92, 15.7%), and BAPS-11 (n= 98, 16.6%) as being the dominant *S. canis* lineages in the whole dataset. In Germany, the dominant *S. canis* clusters are BAPS-5 (n = 143, 33.6% of German isolates) and BAPS-6 (n= 86, 20.1%). We observed that the ST9 isolates (n= 127) clustered within BAPS-5 along with ST3 (n= 18), ST16 (n= 11), ST23 (n= 7), ST29 (n=1), ST48 (n= 1) and unidentified STs (n= 18). This contrasted with BAPS-6 which was composed almost exclusively of ST43 isolates (91/92 isolates) (**Fig 2a**). Given the differences in MLST distribution within BAPS-5 and BAPS-6, we hypothesised that BAPS-6 genomes would have significantly less genetic diversity compared to BAPS-5. To test this, we calculated the pairwise SNP distances of the isolates within each cluster compared to their respective internal references. We found that the pairwise SNP distances of the isolates within BAPS-6 ranged from 10 to 112 SNPs with a median value of 53. This contrasts with the genomes of the BAPS-5 isolates which had pairwise SNP distances ranging from 32 to 886 SNPs with a median value of 787. This confirmed that the BAPS-6 genomes were distinctly less diverse compared to BAPS-5 (**Fig 2b**). Based on their genetic diversity, we hypothesized that BAPS-6 and BAPS-5 isolates would differ in their clinical manifestations. To test this, we determined the proportions of different clinical manifestations exclusively in German dogs. Isolates with unrecorded pathologies were excluded. We found that 71.4% (n= 55/77 German canine isolates) of BAPS-6 isolates were recovered from eye, ear, skin and wound infections whilst this corresponded to 67.9% (n= 57/84 German canine isolates) of the BAPS-5 isolates. We found that one BAPS-5 isolate was recovered from a case of systemic disease whereas BAPS-6 had no cases of systemic infections (**Fig 2c**).

**Fig 2:**
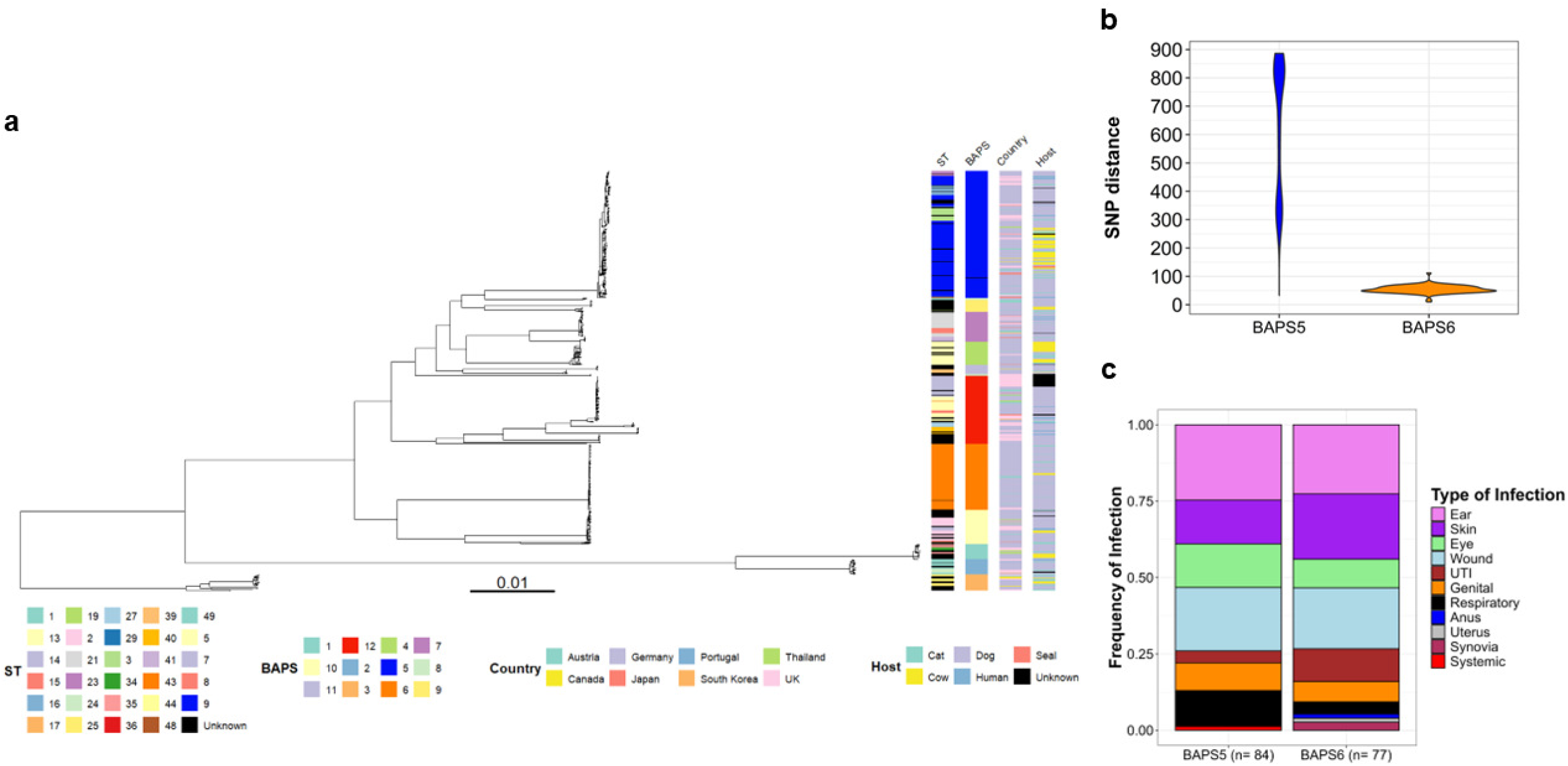
Analysis of the general population structure of *S. canis.* Maximum Likelihood SNPs tree of 585 isolates of *S. canis* rooted to *S. dysgalactiae* and annotated with ST, BAPS cluster, country of origin and host. The tree scale represents the number of substitutions per site (a). Violin plot representing SNP distances of BAPS-5 and BAPS-6 compared to internal references (b). Barplot representing the frequencies of different German canine infections associated with BAPS 5 and BAPS-6 isolates (c).

We then constructed a pan-genome accumulation plot of the entire *S. canis* population to determine whether *S. canis* possessed an open or closed pangenome. No plateau was observed in the pangenome accumulation plot, indicating that *S. canis* has an open pangenome that is receptive to new gene acquisition events **(Fig. S1)**.

### BAPS-6: Expansion through an acquired ICE

To further characterise BAPS-6 isolates in greater depth, we constructed a reference-based maximum likelihood tree to examine how BAPS-6 is internally structured. As was observed in **Fig 2b**, we found that BAPS-6 was comprised of a homogenous population and that the cluster was composed almost exclusively of ST43 isolates from German dogs (**Fig 3a**). We collected samples in both the northern and the southern parts of Germany with the Berlin, Bavarian and North Rhine Westphalian regions having the highest sampling densities (**Fig 1a**). To better understand the expansion of BAPS-6 isolates, we attempted to date its emergence with *BEAST* and found that the most recent common ancestor likely dated to 1988 [95% Highest Posterior Density (HPD): 1961 - 2007] (**Fig 3a**). A major expansion of the BAPS-6 lineage began around 2010 and dominated the population by 2020 **(Fig S2).**

**Fig 3:**
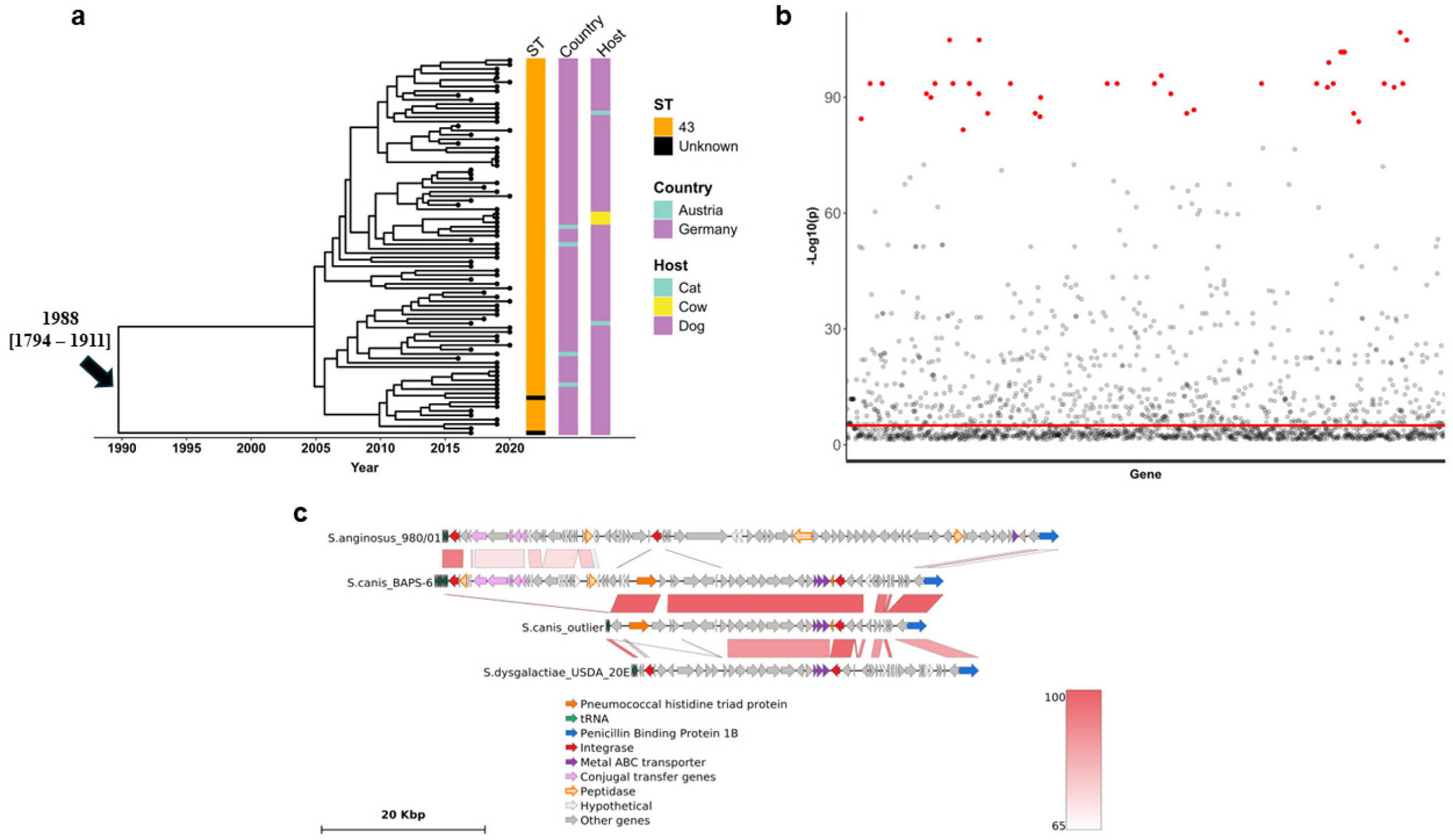
Phylodynamics and GWAS of the rapidly emerging BAPS-6 cluster. Time-scaled Maximum clade credibility (MCC) tree of BAPS-6 cluster associated to its metadata rooted to IMT39221 from the neighbouring BAPS-10 cluster. The X-axis is scaled by year. Scale of tree represents number of substitutions per site and numbers near the root of the tree represent the calculated date of emergence of the cluster with 95% HPD values in brackets (a). Manhattan plot representing all genes significantly associated with BAPS-6. The red line intercepting the Y axis represents the Bonferroni corrected significance threshold set at –Log10(p) > 5. Red dots represent the 49 genes with the highest association to BAPS-6 (b). Genofig representation of the BAPS-6 ICE compared to a similar genetic region in *S. anginosus* (accession: CP118036.1) and a *S. canis* outlier compared to a similar region in *S. dysgalactiae* (CP116779). Genes are annotated based on colours. Red lines linking the different rows indicate nBLAST identity percentages. A darker shade of red indicates a stronger identity. The dotted orange genes in the BAPS-6 ICE represent the putative peptidases (c).

To discover a possible cause for the expansion, we performed a gene presence/absence based GWAS with *Scoary* to identify any overrepresented genes exclusive to the BAPS-6 cluster. We found 49 genes that were significantly associated with the cluster at a –Log10 p-value > 80 and that 27 of these genes were in the same region of the genome (**Fig 3b**). We determined this genetic region to be a putative Integrative and Conjugative Element (ICE) using the *ICEfinder* database. The putative ICE was 20,800 bp long and carries genes for an integrase, four conjugal transfer proteins, one ATP-binding protein as well as two genes corresponding to a toxin-antitoxin system. An *nBLAST* analysis showed that this putative ICE shared more than 65% nucleotide identity with *S. anginosus* (**Fig 3c**). We compared the genomic context of the BAPS-6 ICE with an outlier from the BAPS-12 cluster of *S. canis.* As expected, the ICE was completely absent from this outlier. The ICE inserted between 19 tRNA genes and upstream of a highly conserved region of the *S. canis* genome near the gene for a pneumocococcal-like histidine triad protein. We defined the boundaries of the ICE to be between the tRNA region and the final gene which was not detected in the BAPS-12 outlier (an acyl-coa n-acyltransferase). We also detected three zinc ABC transporter genes, another integrase gene, and the gene for the penicillin binding protein 1B. This conserved region largely resembled the genomic region of *S. dysgalactiae*.

In addition, we identified two genes within the putative ICE as coding for peptidases. One peptidase in the putative ICE represents a putative cysteine protease (C51 family) using the Interpro database. Analysing the protein structure of the C51-family peptidase showed us that the protein possessed seven α-helix structures and one β-sheet structure **(Fig S3a)**. With *PhiGnet*, we found that the C51-family peptidase had two active sites on loop domains with activation scores higher than 0.9 when related to both the *Gene Ontology* (GO) and *Enzyme Classification* (EC) databases **(Fig S3b)**. The first active site was detected to be at the 25^th^ to the 48^th^ amino acid residue. The second active site was around the 100^th^ amino acid residue. We predicted the two active sites to have hydrolase activities according to GO and glycosidase activities according to EC. We identified the second peptidase as an M78-family peptidase again containing seven α-helix structures and one β-sheet structure **(Fig S3c)**. Active sites were significantly less clear with activation scores differing between the GO and EC databases. We found that the peptidase was declared to be involved in metal ion binding by GO and as a metalloendopeptidase by EC **(Fig S3d)**. The regions with activation scores above 0.9 were predicted by GO to be between the 57^th^ and 74^th^ amino acid residue corresponding to a β-sheet domain and the 112^th^ to the 133^rd^ amino acid residue corresponding to α-helix domains. In the EC database, the active sites were between the 116^th^ to the 128^th^ amino acid residue and between the 145^th^ to the 155^th^ amino acid residue, respectively, both corresponding to α-helix domains.

### BAPS-5: Host transition though acquisition of a new *lac* operon

We constructed a reference-based maximum likelihood tree of the BAPS-5 cluster which was put through ancestral state reconstruction with *BEAST.* Twenty-four isolates from the BAPS-5 cluster were removed to adjust for a better temporal signal. We found that BAPS-5 was identified in five different countries (Germany (n=135), UK (n=16), Austria (n=5), Canada (n=2), and Japan (n=1). BAPS-5 isolates also had a wide host range, including dogs (n=107), cows (n=33), cats (n=17), and seals (n=2). We observed clustering of 44/53 non-canine isolates of BAPS-5 into one clade in the tree (**Fig 4a**). We found that these isolates also clustered based on their accessory genome (**Fig 4b**). With time calibrations, we found that BAPS-5 is predicted to have first emerged in a time long before BAPS-6 in the year 1786 (95% HPD: 1645-1840) with a 99.76% probability that it emerged from a canine host. BAPS-5 evolved into three distinct subclusters: two canine-focused clusters which were predicted to have emerged in 1804 (95% HPD: 1674-1853) and 1826 (95% HPD: 1710-1875) as well as one bovine and feline cluster containing 62 isolates in total which was predicted to have emerged in 1879 (95% HPD: 1794-1911). In this bovine-enriched cluster, we identified 20 instances of canine to other host transitions with the oldest transition occurring around 1798 (99% posterior probability of a canine to feline transition, 95% HPD: 1665-1849) and the most recent one in 1993 (89% posterior probability of a canine to feline transition, 95% HPD: 1978-2004). (**Fig 4a**).

**Fig 4:**
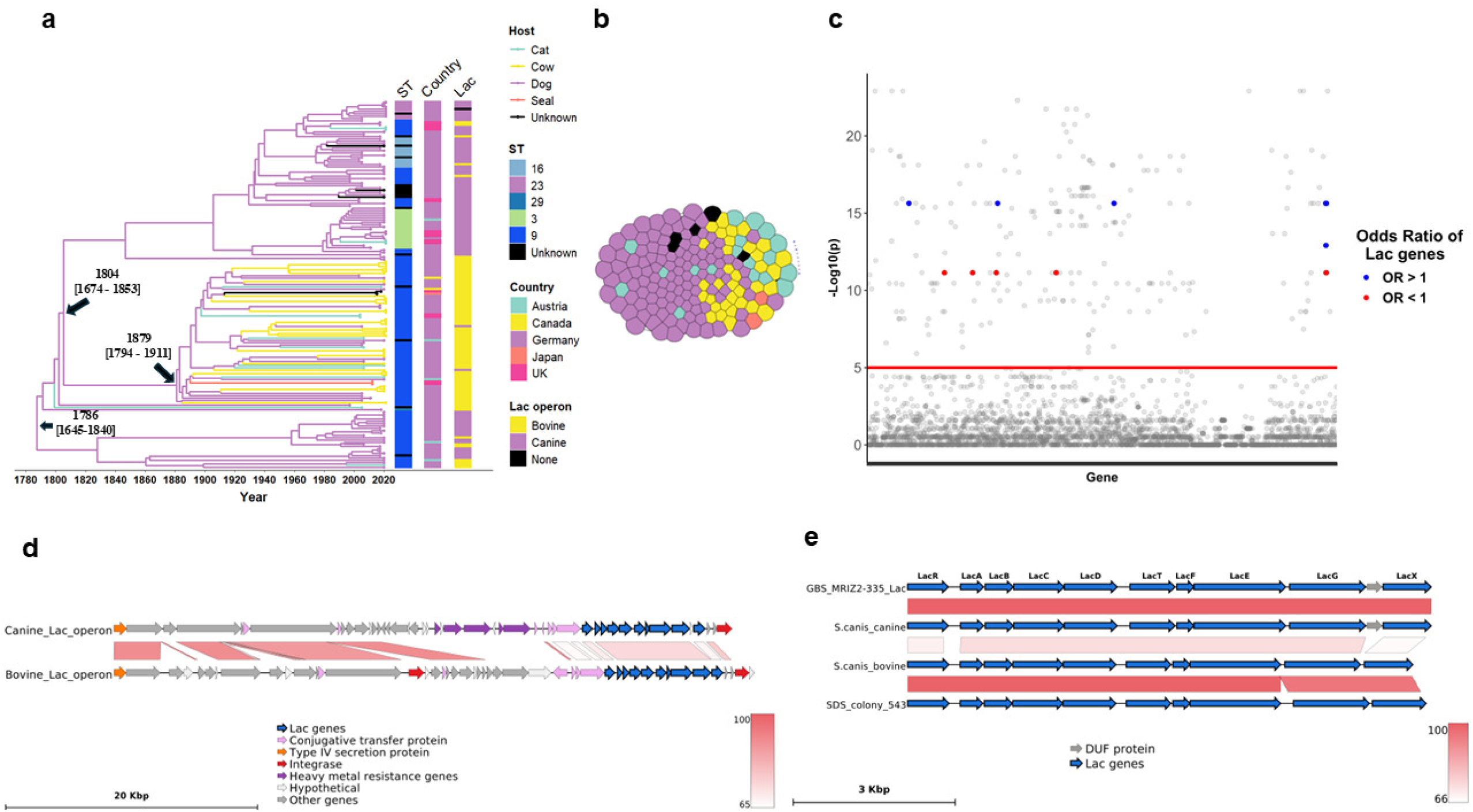
Phylodynamics and GWAS of the multi-host BAPS-5 cluster. Ancestral state reconstruction of the BAPS-5 cluster based on host origin of isolates associated to its metadata. X-axis is a timescale by year and numbers near tree nodes represent the calculated dates of emergence of the main cluster and sub-clusters of BAPS-5 along with 95% HPD values in brackets. The tree is rooted to DRR015857 from the BAPS-4 cluster (a). Accessory genome cluster of BAPS-5 related to host metadata. Host colours are identical to those represented in Fig 4a (b). Manhattan plot of genes significantly associated with the BAPS-5 multi-host isolates. Red line shows cutoff significance of the Bonferroni corrected p-value at –Log10(p) > 5. Blue dots are *lac* genes with Odds Ratios superior to 1 and red dots are *lac* genes with Odds Ratios inferior to 1 (c). Genomic context of the *lac* operon variants in a canine representative of BAPS-5 and a bovine representative of BAPS-5. The genes are coloured based on function. Red lines linking the rows indicate nBLAST identity percentages. A darker shade of red indicates a stronger identity. The *lac* genes are represented in blue (d). Representation of the BAPS-5 canine and bovine/feline *lac* operons compared to the *S. agalactiae* (OX460934) and *S. dysgalactiae* (CP078548) *lac* operons (e).

We conducted a SNPs-based GWAS with *Pyseer* on the core genome of BAPS-5 to identify any mutations that were significantly associated with bovine and feline isolates. We set a threshold for significance with a Bonferroni correction calculated at –Log10(p) > 3.7 and observed that a total of four unique SNPs were associated with these isolates **(Fig. S4a)**. These SNPs occurred in four genes in the core genome of *S. canis*, coding for a CBS-HotDog domain-containing transcription factor (*spx*R), pyruvate oxidase (*spx*B), an aminopeptidase (group_90), and bifunctional (p)ppGpp synthase/hydrolase (*rel*A) **(Table S2)**. We then tested if these genes were undergoing positive selection to see if they could be conferring a fitness advantage to the isolates with *PAML*. The multi-host cluster was determined as the foreground lineage for testing positive selection and the two canine clusters as the background lineage **(Fig. S2)**. We found that overall, these genes showed a median dN/dS of 0.47 showing that they were undergoing purifying selection **(Fig. S4b)**.

The very small number of core genome mutations possibly conferring an advantage to the non-canine isolates led us to analyse the accessory genome for any genes significantly associated with these isolates. We conducted a gene presence/absence-based GWAS with *Scoary* to highlight genes that were significantly associated with bovine and feline isolates. With a Bonferroni correction, we set the threshold for significance at –Log10(p) > 5 and calculated Odds Ratios (OR) with OR > 1 being significantly associated with the canine isolates and OR < 1 being significantly associated with the bovine and feline isolates (**Fig. 4c**). We found that a total of 223 genes showed significant associations with one of these groups. Notably, there was a reciprocal association with two alternative *lac* operons: Bovine and feline isolates were significantly associated with one *lac* operon variant whereas the canine isolates were associated with a different *lac* operon. Both of these *lac* operons were distinct from the one identified in the core genome of BAPS-5. We observed clustering of the alternative *lac* operons on the tree based on host origin (**Fig 4a**). We analysed the genomic context of the lac operons and found that both were located upstream of an integrase gene and downstream of several conjugative transfer genes and a type IV secretion system gene in the case of the bovine isolate (**Fig 4d**). We found several heavy metal-associated genes corresponding to copper and cadmium resistance upstream of the canine lac operon but absent in the region upstream of the bovine lac operon. The bovine lac operon instead possessed several genes for ATP-dependent proteins. The rest of the genomic context was significantly conserved in both canine and bovine isolates. With *nBLAST*, we found that the canine *lac* operon was near-identical to an *S. agalactiae lac* operon, whereas the bovine *lac* operon was near-identical to an *S. dysgalactiae lac* operon. However, when we compared the two *S. canis lac* operons, we found that they only shared approximately 65% nucleotide sequence identity and that the canine *lac* operon had an additional gene for a protein with a domain of unknown function (DUF) immediately upstream of the *lacX* gene. (**Fig 4e**).

## Discussion

We have produced the largest and highest resolution *S. canis* population genetics study to date providing a more representative view of the population structure of this important canine pathogen. With 585 *S. canis* isolates of isolated between 1996 and 2020, we describe two dominant BAPS clusters associated with German canine hosts: the BAPS-5 cluster which predominated from 1996 to 2015 and the BAPS-6 cluster which increased in prevalence from 2016 to 2020. We found that BAPS-6 is a clonal subpopulation almost exclusively associated with the presence of an ICE which most likely was acquired through genetic exchange with *S. anginosus*. This ICE contains a putative cysteine protease of the C51 family, a potential virulence factor that is often associated with streptococci and a putative M78-family peptidase. We found that BAPS-5 is an older and more diverse cluster of likely canine origin which first emerged around the year 1786 and spread into bovine and feline hosts from 1879 onwards. The bovine and feline isolates of the BAPS-5 cluster have a unique *lac* operon with a high level of nucleotide sequence identity to a similar operon in *S. dysgalactiae.* These genes may confer an advantage to *S. canis* when infecting feline and bovine hosts.

The prevalence of the BAPS-5/ST9 population of *S. canis* inside and outside of Germany is unsurprising as this has already been recorded in Germany and Portugal in previous studies (2, 11, 27). However, the sudden rise of the BAPS-6/ST43 population of *S. canis* is an unexpected observation due to it either being absent from previous datasets or being underreported (2, 11, 27). More interesting is the detected shift in dominance within Germany, but not in other parts of the world suggesting that this might for now be a contained event. The sudden detection of BAPS-6 and the recent emergence of the cluster at the year 1988 coincide with its lower genetic diversity since there would have been little time for significant genetic divergence. The BAPS-6 and BAPS-5 cluster appear to have a similar tendency to cause eye, ear, and skin infections in dogs. We did however observe that in this dataset, BAPS-6 isolates have yet to be recorded to cause systemic infections unlike BAPS-5 although this might be due to the higher sampling density of BAPS-5 rather than any difference in disease manifestation.

The almost exclusive and universal presence of the ICE in BAPS-6 indicates a possible reason for its sudden expansion in Germany. A similar phenomenon has been observed in the human pathogen *S. pyogenes* where several ICEs containing multi-drug resistance and toxin markers were significantly associated with the emergence of an epidemic of scarlet fever in Hong Kong (28). The nearest relative to *S. canis*, *S. dysgalactiae*, has also shown the capacity to suddenly expand due to the acquisition of an ICE containing the *erm*(B) and *tet*(M) drug resistance genes. This was recorded in cases of bacteremia in the Kyoto-Shiga region of Japan between the years 2005 and 2021 (30). The BAPS-6 ICE appears to have the most similarity with a genetic region in *S. anginosus,* an emerging human gastro-intestinal pathogen (32). To date, there has only been one recording of the presence of *S. anginosus* on a canine host where it was associated with a human case of infection caused by a dog bite (33). The homology of the BAPS-6 ICE with *S. anginosus* suggests that a genetic exchange occurred between the two, but the overall 65% nucleotide identity is relatively low. This indicates that the clade of *S. anginosus* represented by the genome sequence is probably not the immediate donor, and that this may be an unsampled clade of *S. anginosus* or another streptococcal species. Given the lack of sequences with a higher identity to the ICE in the entire GenBank database, it is difficult to clarify its immediate source.

When analysing the M78-family peptidase in the ICE, the predicted functions as metal ion binding and as a metallo endopeptidase coincide with its attribution to the M78-family. M78-family peptidases have been shown to bind zinc though their HEXXH motif allowing for proteolysis (34). The genomic context of the ICE showed that it had inserted upstream of several zinc-related ABC transporter genes and the gene for the zinc-binding pneumococcal histidine triad protein (35, 36). It is possible that the zinc bound by the zinc ABC transporters and the pneumococcal histidine triad protein could facilitate zinc availability for the M78-family peptidase and encourage its activity. M78 peptidases are known to cleave the ImmR protein responsible for repressing horizontal gene transfer of mobile elements (34). The ImmR protein is, however, absent in this ICE which suggests that the M78 peptidase might have another function in this context encouraged by the increased zinc availability via the proteins encoded by the genes downstream of it.

The C51-family peptidase present within the ICE could be an interesting potential virulence factor for *S. canis.* Cysteine proteases are strongly related to the pathogenesis of *S. pyogenes* with IdeS and SpeB being the most well-characterised proteins in this category (37). These cysteine proteases have shown strong IgG degrading capabilities which would aid in host immune evasion during infection (37). IdeS and SpeB have also shown the capacity to bind integrins through the RGD motif which would allow the bacterium to invade host cells through host-dependent internalization (38, 39). IdeS appears to be more restricted in its function with its only other recorded binding capabilities being for IgA and IgM, further contributing to the immune evasion strategies of *S. pyogenes* (40). SpeB appears to be a much more versatile protein capable of degrading a variety of host proteins, such as all the immunoglobulin classes and C3b which would aid in host immune evasion (41). It can also degrade fibrinogen, plasminogen, vitronectin, and fibronectin which could facilitate invasion and spread in the host (42–44). Moreover, SpeB shows activities directly related to proteins and systems of *S. pyogenes* itself. It can cleave the M protein at the N-terminal or C-terminal site allowing its dissociation from the bacterial cell surface which could alter the protein binding capabilities of *S. pyogenes* (45). It has been shown to degrade other exotoxins, such as streptokinase which would regulate its virulence (46). This gives many directions for research concerning the function of the BAPS-6 C51 peptidase. Further understanding the pathogenesis of the BAPS-6 isolates could become vital in the veterinary field. A study of *S. canis* canine infections in the United States indicated that ST43/BAPS-6 was associated with treatment failure of corneal ulcerations (47). The continued rise of BAPS-6 could be followed by increased treatment failure of these types of infection.

The age of the BAPS-5 cluster of *S. canis* could be the main reason it has dominated the entire population of *S. canis* in previous population genetics studies (2, 11, 27). The earlier emergence and wider spread of the cluster could have allowed it more time and opportunity to adapt to the observed additional bovine and feline hosts. This could also be an explanation for the strong presence of the cluster outside of Germany. The emergence of the bovine/feline subcluster in Germany around the year 1879 coincides with the beginning of the dairy industry around the year 1870 (48). The dairy industry expanded shortly thereafter with dairy consumption rising from 284 kg/capita in 1860 to 355 kg/capita in 1900 (48). The consequent increase in cattle population to meet demands would have encouraged the transition of *S. canis* to the bovine host. The strong presence of feline isolates in the cluster suggests that there is cross-infection of feline and bovine hosts by *S. canis* in these farms as shown previously, although in this study we did not find host transitions of this type with ancestral state reconstruction (14). Instead, BAPS-5 appears to transition primarily from dog to cow or cat which encourages the possibility that canine hosts are a major source of transmission to bovine and feline hosts. When searching for host-associated genetic signatures, we observed purifying selection of all core genes with mutations significantly associated with the non-canine isolates. This is unsurprising since most mutations are thought to be deleterious (49). This suggests that the bovine/feline sub-clade did not likely emerge due to advantageous mutations in the core genome. Significant differences were found in the accessory genome of the non-canine isolates compared to the canine isolates. Notably, the finding of the reciprocal canine and non-canine *lac* operon variants implies that certain *lac* operon variants are advantageous depending on the host context. *S. agalactiae* is known to have a variant of the *lac* operon linked to bovine host adaptation: Lac.2 (26, 50, 51). The same *lac* operon was detected in another bovine pathogen, *S. dysgalactiae* subsp. *dysgalactiae* (52). Here, we present evidence of possible *S. canis* host adaptation to bovine and feline hosts.

The BAPS-5 canine *lac* operon was located next to four genes involved in copper and cadmium resistance. Heavy metals are highly abundant in the environment due to pollution of soil and water by industrial and agricultural practices (53). This contributes to the selection of any genes that might be neighbouring the heavy metal resistance genes (53, 54). Notably, the presence of heavy metal resistance genes is often associated with co-selection for antimicrobial resistance which contributes to the growing threat of multi-resistant bacteria (55). Furthermore, it has been shown that heavy metal resistance could be directly related to mobile elements containing virulence factors for *Pseudomonas aeruginosa* and, therefore, that heavy metal resistance could be associated with increased pathogenesis of this bacteria (56). The presence of the heavy metal resistance genes in the canine isolates rather than the bovine isolates could be because of the increased bioaccumulation of heavy metals in animals which are in a higher position in the food chain (57). Indeed, it has been shown that pet feed contains higher than tolerable quantities of heavy metals such as, cadmium, lead, and arsenic (58). This could provide a source for the selection of heavy metal resistance genes in canine hosts. The selection for the *lac* operon in bovine hosts might purely be due to the advantage it could confer to *S. canis* in the presence of milk as has been observed for *S. agalactiae* (50). The canine *lac* operon has an added DUF protein immediately upstream of the *lacX* gene which codes for an aldolase 1-epimerase (51). This enzyme catalyzes the first step in galactose metabolism by converting α-D-galactose into β-D-galactose. The presence of the DUF protein in the canine isolates could affect the expression of *lacX*.

Our study provides detailed information on *S. canis* expansion in Germany. It would be interesting to expand these findings by adding more isolates from different countries and hosts to evaluate whether we see differences in lineage dominance depending on these factors. *S. canis* is often associated with canine infections, but is equally prevalent in feline hosts (59). Our dataset lacks a large number of feline isolates and could benefit from future analyses that target the evolution of *S. canis* in this context. The groundwork for potential phylogeographic studies has also been laid. An *S. canis* study was conducted in the United Kingdom where the zoonotic potential of the pathogen was elucidated (2). BAPS-6/ST43 isolates have been detected in the United States and the relevance of the BAPS-5 population was first emphasized by two studies executed in Portugal where ST9 in both cases was shown to be the dominant population of *S. canis* (11, 27, 47). Including more isolates from these countries could strengthen the dataset sufficiently to perform phylogeographic analyses and infer any exchanges between the locales. One final limitation of this study is the lack of experimental evidence pertaining to the ICE of BAPS-6 and the *lac* operon of the non-canine BAPS-5 isolates. To determine whether the ICE confers an advantage to BAPS-6 in the context of infection, the functionality of the C51-family peptidase contained within it must first be characterised experimentally. IgG cleavage could be a possible candidate which would contribute to *S. canis* immune evasion although as mentioned previously, cysteine proteases have been recorded to have many different functions which equally warrant more studies. In the case of the newly identified *lac* operon of BAPS-5, a comparative analysis of the non-canine and canine isolates should be conducted with lactose degradation assays to measure whether the operon has an impact.

In conclusion, this study is the largest population genomics analysis for *S. canis* to date. For the first time, through phylodynamics and GWAS, we have associated the rapid expansion of a pathogenic lineage of *S. canis* with the presence of a putative ICE containing a putative cysteine protease. This is also the first study to link the host transition of *S. canis* from canine to bovine/feline hosts to the acquisition of a variant of the *lac* operon. Overall, we highlight the need to carefully examine *S. canis* as a major veterinary threat in Germany for canine, feline, and bovine hosts alike.

## Material and Methods

### Strain selection and sequencing

*Streptococcus canis* strains were selected by first cultivating the isolates on Columbia blood agar plates with 5% sheep blood at 37°C overnight. Any strains that were shown to be β-haemolytic were then checked for Lancefield type G expression with the PathoDxtra™ Strep Grouping Kit (Thermo Fisher Scientific, Waltham, MA USA). DNA was extracted from Lancefield G-positive strains with the QIAamp DNA Mini Kit (QIAGEN, Venlo, Netherlands) following the standard Gram-positive protocol. To identify whether the strains belonged to the *S. canis* species, Lancefield type G strains were screened via Maldi-TOF. The confirmed *S. canis* strains were prepared for sequencing using the Nextera XT DNA Library Preparation Kit (Illumina, Inc., San Diego, CA, USA) according to the manufacturer’s recommendations, followed by 2 × 300-bp paired-end sequencing in 40-fold multiplexes on the MiSeq platform with the MiSeq reagent kit v3 (Illumina, Inc.).

### Sequence analysis and assembly

FastQ files were quality assessed with *FastQC* v0.12.1. Sequence reads were classified using *Kraken* v0.10.6 to confirm classification to the *Streptococcus* genus. The raw reads were passed through the *assembleBAC* pipeline for assembly via the integrated *Shovill* pipeline, MLST classification with the integrated *mlst* v2.23.0 program and annotation with the integrated *Bakta* v1.9.2 program (https://github.com/avantonder/assembleBAC). The raw reads were then put through the *Bactmap* pipeline for quality and adapter trimming via the integrated *fastp* v0.23.4 program (60). The entire dataset was then mapped to the HL_77_2 *Streptococcus canis* reference genome [NCBI RefSeq assembly GCF_010993845.2]. *Gubbins* v3.3 was run on the mapped assemblies to remove loci associated with recombination events (61). Internal mapping was conducted on BAPS-5 and BAPS-6 using the same methodology as described for the whole dataset. The reference for BAPS-5 was AGF1071 and the reference for BAPS-6 was AGF1279. Genomic regions of interest were visualized and compared with *Genofig* (62). Protein structure and active sites were determined with the PhiGnet webtool (63).

### Phylogenetics and phylodynamics

Maximum Likelihood phylogenetic trees were constructed from reference-based alignments produced after masking recombinant regions. *IQtree* v2.0.3 was run on these alignments, accounting for constant sites, using the best fit model (MODEL) and 1000 bootstrap replicates (64–67). Pairwise SNP distances were inferred with the *pairsnp* R package (https://github.com/gtonkinhill/pairsnp). Temporal signals were evaluated with *TempEst* v1.5.3 (68). *BEAST* v1.10.4 was used to date the trees constructed previously by using a GTR substitution model, uncorrelated relaxed clock and a coalescent constant size model (69). The discrete trait in BAPS-5 was defined as the host origin of the isolates.

### Pangenome analysis

#### Genome wide association study (GWAS)

To calculate a pan-genome for *S. canis*, *Panaroo* v1.3.4 was run on the entire dataset in strict clean mode while defining the core threshold at 0.95 (70). The resulting gene presence/absence file was used to run *Scoary* v1.6.14 at default settings (71). BAPS-6 was defined as the phenotype for the analysis and significance was defined as Bonferroni p-value < 0.05 and specificity > 90%. GWAS on the BAPS-5 cluster was conducted on its own core genome inferred with *Panaroo* v1.3.4. *Pyseer* v.1.3.10 was used while using the SNPs of the core genome as the association model (72–74). The phenotype was defined as any isolate that is not of canine origin and significance was determined with an LRT p-value.

#### Accessory genome clustering

The accessory genome was defined as any genes present in less than 95% of the isolates. The pairwise distance matrix of the accessory genes was produced by *GraPPLE* (75). The accessory genome was visualized on *Graphia* (76).

#### Pangenome accumulation plot

The pangenome accumulation plot was produced with the gene presence/absence matrix produced by *Panaroo* v1.3.4 coupled with the *Panstripe* R package (77).

### Gene selection analysis

Gene selection analysis was conducted with the *PAML* package (78). *CODEML* was run on select genes in the core genome of the *S. canis* BAPS-5 cluster with the branch model. A maximum likelihood tree of the core genome was generated with *IQtree* v2.0.3 with the same parameters as described for the mapped genome trees. The foreground lineage was defined as the bovine and feline cluster. The background lineage was defined as the canine cluster. dN/dS values were then extracted from the resulting analysis.

## Acknowledgements

We would like to thank Julian Brombach (Institute of Microbiology and Epizootics, Freie Universität Berlin) for excellent technical assistance. We are grateful to Thomas Peters (Milchtierherden-Betreuungs-und Forschungsgesellschaft mbH, MBFG, Wunstorf, Germany), Julie-Hélène Fairbrother (Laboratoire de santé animale de Saint-Hyacinthe, Ministère de l’Agriculture, des Pêcheries et de l’Alimentation du Québec (MAPAQ), Saint-Hyacinthe, Québec, Canada), Reglindis Huber (Eutergesundheitsdienst – Tiergesundheitsdienst Bayern e.V., Poing, Germany), John Timoney (Department of Veterinary Science, University of Kentucky, Lexington, US), Georg Wolf (Institute for Infectious Diseases and Zoonoses, Department of Veterinary Sciences, Faculty of Veterinary Medicine LMU Munich, Germany), Jörg Rau (Chemisches und Veterinäruntersuchungsamt Stuttgart, Germany), and Maria Kordesch (InVitro, Wien, Austria) for providing *S. canis* strains. This work was supported by the Petplan Charitable Trust foundation (grant no. S20-851-890 to EA, JP, and MF), as well as by the German Research Foundation (DFG, grant no. FU 1027/3-1 to MF and FU 1027/5-1 to MMDK and MF).

## Data Availability

Newly sequenced isolates were submitted to the Sequence Read Archive as fastq files under Bioproject number PRJNA1332248.

## Author Contributions

Conceptualization – EA, AJvT, AW, JP, MF

Data Curation – EA, MMDK, AML, ALB

Formal analysis – EA, MMDK

Funding Acquisition – AW, JP, MF

Investigation – EA, AML, ALB, MF

Methodology – AjvT, ALB

Project Administration – JP, MF

Resources – TS, SS, ALB, JP, MF

Software – TS, JP

Supervision – AjvT, HJW, AW, JP, MF

Validation – MMDK, HJW, TS, ALB, JP, MF

Visualization – EA, AJvT

Writing - Original Draft Preparation – EA, MMDK, AML, ALB

Writing - Review and Editing – AjvT, HJW, SS, AW, JP, MF

## Supporting Information

**S1 Fig: Pangenome accumulation plot of the entire *S. canis* dataset.** Calculated based on accessory genome size and the number of included genomes.

**S2 Fig: Bayesian skyline plot of the BAPS-6 cluster.** Represented by effective sampling size on the Y-axis and time in years on X-axis.

**S3 Fig: Protein structure and functional analysis of the C51 and M78 peptidases.** Blue shades on the protein represent α-helix structures and green shades indicated β-barrel structures (a, c). Biological activation scores calculated at each residue of the C51 peptidase based on Enzyme Classification (top plot) and Gene Ontology (bottom plot) (b, d).

**S4 Fig: Analysis of host bovine host adapatation in the core genome.** Manhattan plot of SNPs significantly associated with the bovine BAPS-5 isolates (a). Red line shows cutoff significance of the Bonferroni corrected p-value at –Log10(p) > 3.72. Boxplot representing the overall dN/dS ratio of the genes with SNPs significantly associated with the bovine strains (c). Red line represents threshold for asserting positive selection (dN/dS > 1).

## S1 Table: Dataset metadata

**S1 Table: Dataset metadata.**

**S2 Table: Table representing the dN/dS values of the core genes with SNPs. significantly associated to the multi-host BAPS-5 isolates.**

**S3 Table: Scoary results of BAPS-6 isolates in comparison to whole dataset . S4 Table: Scoary results of non-canine isolates in BAPS-5.**

